# Synthetic Histology Images for Training AI Models: A Novel Approach to Improve Prostate Cancer Diagnosis

**DOI:** 10.1101/2024.01.25.577225

**Authors:** Derek J Van Booven, Cheng-Bang Chen, Oleksander Kryvenko, Sanoj Punnen, Victor Sandoval, Sheetal Malpani, Ahmed Noman, Farhan Ismael, Andres Briseño, Yujie Wang, Himanshu Arora

**Author notes:** **Correspondence:** Himanshu Arora.

## Abstract

Prostate cancer (PCa) poses significant challenges for timely diagnosis and prognosis, leading to high mortality rates and increased disease risk and treatment costs. Recent advancements in machine learning and digital imagery offer promising potential for developing automated and objective assessment pipelines that can reduce human capital and resource costs. However, the reliance of AI models on large amounts of clinical data for training presents a significant challenge, as this data is often biased, lacking diversity, and not readily available. Here we aim to address this limitation by employing customized generative adversarial network (GAN) models to produce high-quality synthetic images of different PCa grades (radical prostatectomy (RP)) and needle biopsies, which were customized to account for the granularity associated with each Gleason grade. The generated images were subjected to multiple rounds of benchmarking, quantifications and quality control assessment before being used to train an AI model (EfficientNet) for grading digital histology images of adenocarcinoma specimens (RP sections) and needle biopsies obtained from the PANDA challenge repository. Validation was performed using the AI model trained with synthetic data to grade digital histology from the cancer genome atlas (TCGA) (RP sections) and needle biopsy data from Radboud University Medical Center and Karolinska Institute. Results demonstrated that the AI model trained with a combination of image patches derived from original and enhanced synthetic images outperformed the model trained with original digital histology images. Together, this study demonstrates the potential of customized GAN models to generate a large cohort of synthetic data that can train AI models to effectively grade PCa specimens. This approach could potentially eliminate the need for extensive clinical data for training any AI model in the domain of digital imagery, leading to cost and time-effective diagnosis and prognosis.

---

Prostate cancer (PCa) is the third most commonly diagnosed cancer worldwide and the fifth leading cause of cancer-related death among males ^1–7^. The diagnostic pipeline for PCa begins with measuring prostate-specific antigen (PSA) levels ^8–19^. If the PSA levels are above the normal range, the patient is evaluated using one of several assays such as 4Kscore ^20–26^, PCA3 ^27–33^, or phi ^34–39^. Positive results on these assays may prompt imaging studies, such as MRI, to identify potential areas of PCa ^40–45^. Clinicians may also perform biopsies for further evaluation by pathologists and genomic testing ^46–50^. This diagnostic pathway is commonly used to assess PCa prognosis. Multiple follow-up appointments are recommended, especially for men under active surveillance ^47–50^. Notably, over 77% of men with localized PCa are suitable for active surveillance, and this number has increased from 15.5% to 42.2% over the last decade. These men require repeat follow-ups every six months or 1 year if enrolled ^51–53^. However, overdiagnosis in PCa contributes significantly to physical, mental, and sexual health problems ^54^. Therefore, there is a clear need for more effective diagnostic methods for PCa to improve patient outcomes.

In recent years, there has been notable progress in employing machine learning to automate various stages of PCa diagnosis and prognosis ^55^. Machine learning algorithms can learn from data and derive predictions or decisions based on that learning ^55–61^. In the context of PCa, these algorithms can be trained on data from tissue samples, medical images, and other clinical information to identify patterns and features associated with the disease ^62–72^. Machine learning has proven to be particularly effective in the automated analysis of medical images, including MRI scans and biopsy slides ^73–82^. These algorithms can accurately detect suspicious areas of the prostate and provide a more precise diagnosis than traditional manual analysis. PathAI, HTL Limited, PaigeAI, and Deciplex are some of the technologies that leverage digital imagery to assist in diagnosis. These technologies convert digital histology into meaningful data, which could accurately differentiate cancerous and non-cancerous areas ^83–87^.

In spite of these strengths, the available technologies for proactive adaptation in clinics are limited due to the available tools’ ineffective incorporation of cancer heterogeneity. This is attributed mainly to the use of training data, which is primarily derived from clinical trials ^88–91^. There are several limitations to using clinical trial data in digital pathology for training machine learning models ^85,92^. For example, one major limitation is the potential for bias in the data, as clinical trial data is usually collected from a specific group of patients that may not be representative of the general population or a particular stage of disease ^93–95^. This can lead to learning models performing well on the training data but poorly on new or unseen data. Additionally, issues with the quality and reliability of the data may arise due to various factors such as staining techniques, slide preparation, and the microscope’s resolution, which can introduce noise and variability into the data, impacting the performance of machine learning models ^96,97^. Moreover, there may be ethical concerns surrounding the use of clinical trial data in digital pathology, mainly as the data may contain sensitive information about patients, such as their medical history and personal information ^98^. It is, therefore, essential to protect and use this information in accordance with ethical guidelines to ensure the privacy and dignity of patients To address the limitations, one potential solution is to reduce reliance on clinical trial data for training AI models. In this study, we present a novel approach that employs customized generative adversarial network (GAN) models to generate synthetic data for digital pathology applications, using a limited number of original images from radical prostatectomy patients and needle biopsies at various cancer stages. We then utilized the image patches derived from enhanced synthetic data alongside patches from original images to train an AI model (EfficientNet) for grading digital histology sections. Simultaneously, we also trained the AI model using only image patches derived from original digital histology images to grade the same sections. Notably, the AI model trained with synthetically generated data demonstrated superior performance when grading the sections compared to the model trained solely with original data. To validate the results, we employed radical prostatectomy sections from diverse sources, including the Cancer Genome Atlas (TCGA), the University of Miami pathology core, and needle biopsy data from Radboud University Medical Center and the Karolinska Institute (the PANDA challenge)^99^. Our study represents a significant advancement in developing efficient diagnostic and prognostic machine learning models within the realm of digital imagery. By employing innovative conceptual and technical approaches, we successfully overcome a major challenge related to data scarcity in training AI models for digital pathology.

## Network Architecture and Model Selection

To identify the most suitable CNN model for our study, we first evaluated the AlexNet network architecture. This was the first large-scale convolutional neural network to be defined, featuring a series of convolutional pooling layers and an interconnected output layer. To test the accuracy of this model, we randomly selected 20 histology images from the Prostate Adenocarcinoma dataset (obtained from TCGA), which included images from 500 patients, with 4 images from each of the following score: normal, GS6, GS7, GS8, and GS9. We graded these images using default parameters without pretraining the model and compared the results against TCGA scoring and the assessments of two independent pathologists (considered the gold standard).The AlexNet model demonstrated an accuracy rate of 55% in alignment with the gold standard. Next, we evaluated the ResNet model. ResNet incorporates residual networks based on skip connections within the network, which significantly improves upon AlexNet. We tested ResNet across the same 20 images and found that it too, achieved an accuracy rate of approximately 55% in alignment with the gold standard. We then assessed the Xception model. Based on the Inception model, this improves traditional network architectures such as AlexNet. Traditional models attempt to evaluate space and depth in their convolutional layers, whereas Xception separates the two into different dimensions. Xception achieved an accuracy rate of 60% in alignment with the gold standard, outperforming the other two models. Lastly, we tested the EfficientNet model. EfficientNet utilizes a novel scaling method that employs a compound coefficient to scale up a CNN model in a more structured manner. This improves the model’s performance and efficiency while requiring minimal layers to train on existing images. EfficientNet achieved an accuracy rate of 65% in alignment with the gold standard, outperforming the other three models. Based on these results, we have selected the EfficientNet model for our study (Supplementary Figure 1 and Supplementary Table 1).

## Image Preprocessing

Several factors, such as staining protocols, tissue quality, section thickness, tissue folding, and the amount of tissue on the slide, could negatively impact the accuracy of AI models in making predictions from tissue biopsy images. To account for this, we conducted pre-processing normalization of the TCGA images. Specifically, we selected all 500 images and evaluated their color distribution by calculating the mean value of RGB colors and normalizing this value. Images with an RGB mean intensity value that was 2 standard deviations away from the total mean value of all samples were identified as outliers and removed from the dataset. In total, 21 images were discarded due to being outliers. Similar pre-processing was also conducted for the RP section images obtained from University of Miami Pathology core (n=32), where the mean intensity value was calculated and compared to each image. One sample was identified as an outlier and removed from the dataset. For the normalization of needle biopsy slides from Radbound University Medical Center and Karolinska Institute (n=3949), the images were first evaluated for areas of tissue before subjecting them to color normalization. Normalization led to the exclusion of 257 images from the dataset. Overall, our pre-processing steps helped to reduce the variability in tissue biopsy images and ensured a more consistent dataset for the AI models to make predictions.

## Generating Synthetic Images from Prostate Cancer

RP sections (from TCGA and UM) were separated into pseudo-cohorts based on the primary and secondary Gleason pattern and were inspected using the HistoQC software to remove samples of low quality. A random selection of 143 sections was then provided to two pathologists for Gleason pattern annotation. To build our gold standard training data, we required both pathologists to agree with each other and TCGA scoring, as well as precise overlapping annotation of the tumor region. This resulted in a total of 33 images. Image patches from these 25 individuals from TCGA and 8 images from in-house selection were sliced using PyHIST software with dimensions of 96×96 and 256×256 pixel lengths. The areas of annotation obtained from the pathologists were overlaid with these sections, and image patches were required to contain at least 75% tissue image and no more than 25% whitespace. The smallest area of overlap between pathologists was 96 pixels, which was considered the smallest size for image analysis. In total, 219 patches were generated from 33 images, which were enhanced by augmentation to create a total of 2082 patches (Figure 1A).

**Figure 1.**
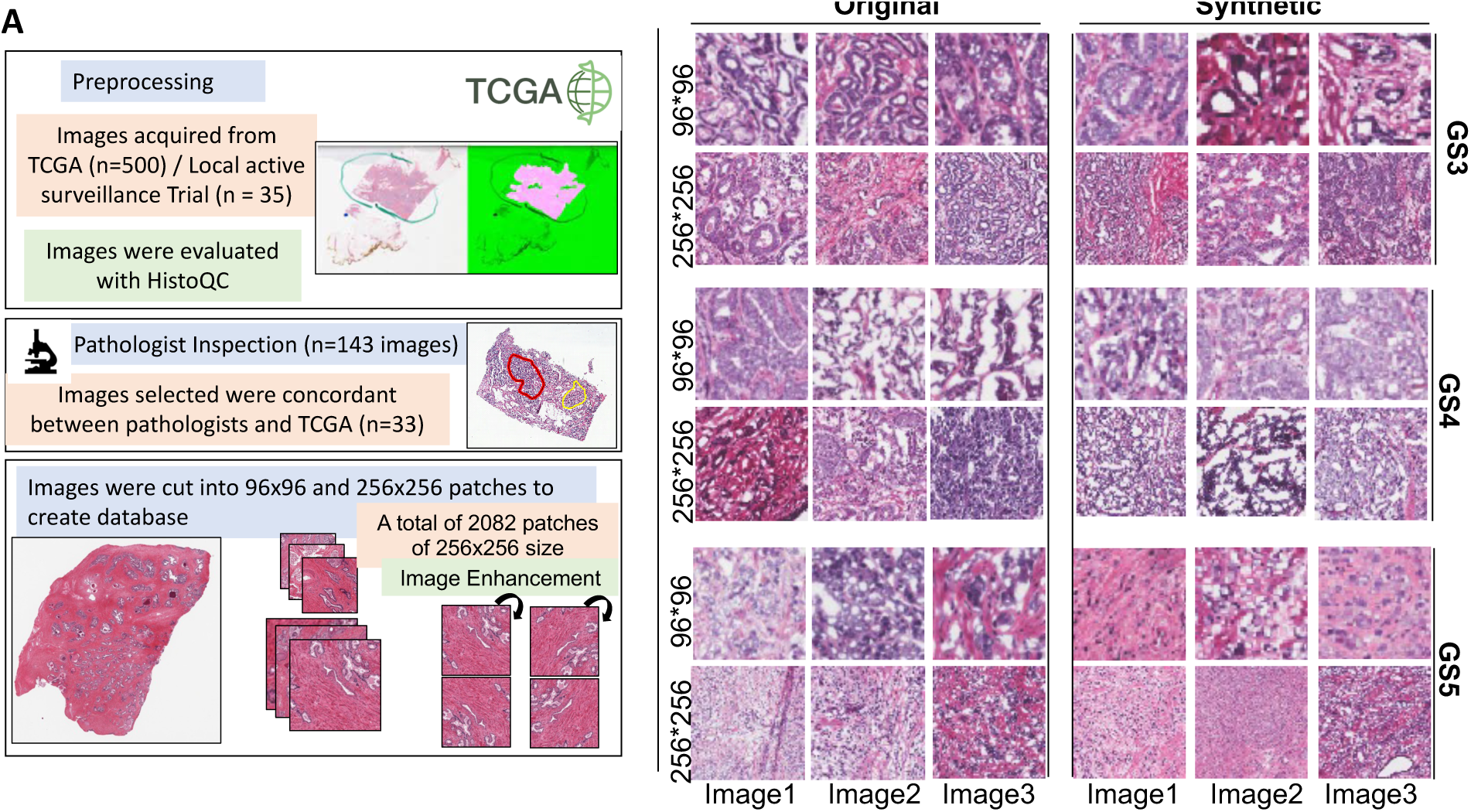
(A) Illustration showing pipeline used in generating synthetic images from prostate cancer digital histology. Images were preprocessed by PyHist and HistoQC. Those that passed QC were then given to pathologist for scoring, and then cut into small patches for modeling. (B) Original and Synthetic images were generated for each primary Gleason pattern 3, 4, and 5, respectively.

## GAN Model Selection

To evaluate the performance of various GAN architectures and select the most appropriate one, 2082 RP image patches extracted from 33 individuals were divided into training cohorts according to their Gleason pattern (GS3, GS4, or GS5). Each training cohort was subsequently subjected to cGAN, StyleGAN, and dcGAN architectures. A total of 1000 synthetic images generated by each GAN were fed into a generic Convolutional Neural Network (CNN) for classification into their respective Gleason categories. The cGAN achieved an accuracy of 0.59, while the StyleGAN and dcGAN demonstrated accuracies of 0.65 and 0.64, respectively. Although the StyleGAN and dcGAN exhibited similar accuracies, their execution times differed significantly when utilizing a standard NVIDIA T100 GPU processor. The generation of 1,000 images in the StyleGAN required 2,372 minutes, whereas the dcGAN completed the task in 901 minutes. Based on these findings, the deep convolutional GAN (dcGAN) network generator was selected and used to create random synthetic images from 2082 patches to create an effective training dataset.

## Benchmarking and. Synthetic Image Patch Synthesis

The resulting image patches from dcGAN were subjected to the Adam optimization algorithm to determine the optimal iteration value, and an optimal iteration of 14,000 was selected (Supplementary Figure 2). To ensure that the correct patch size was taken, random synthetic prostate images generated from the 128×128 and 256×256 pixel patches were subjected to model-based QC assessment. Manual board-certified pathologist inspection of these images was centered around the sharpness of the image and resolution (Figure 1B). The pathologists’ response indicated an 80% approval rate for the data to be sufficiently good for both patch sizes. To further analyze the quality control (QC), we created a simple classification scalable vector machine model where synthetic images were classified into the correct Gleason pattern. Furthermore, the synthetic images generated from the GAN were subjected to index calculations to evaluate the similarity between the images, computed via the relative inception score (RIS). The inception distance, which measures the similarity between images, was also calculated and found to be 17.2 +/-0.15, well above the threshold of 5 typically used in facial recognition GANs, suggesting that the synthetic images are of sufficient quality.

Similarly, we obtained 599 digital pathology images for the prostate from the GTEx Portal, a repository of 25,713 images, and subjected them to color normalization. After correction and normalization, 572 images were used to train the dc-GAN model. The average time taken for a typical GAN run was benchmarked at 2.5 hours per run on a single NVIDIA A1000 GPU chip. Each dc-GAN run yielded 1000 synthetic images that were selected when the generator and discriminator loss functions were equal, which occurred between 25,544 to 62,426 epochs (Figure 2 and Supplementary Figure 2).

**Figure 2.**
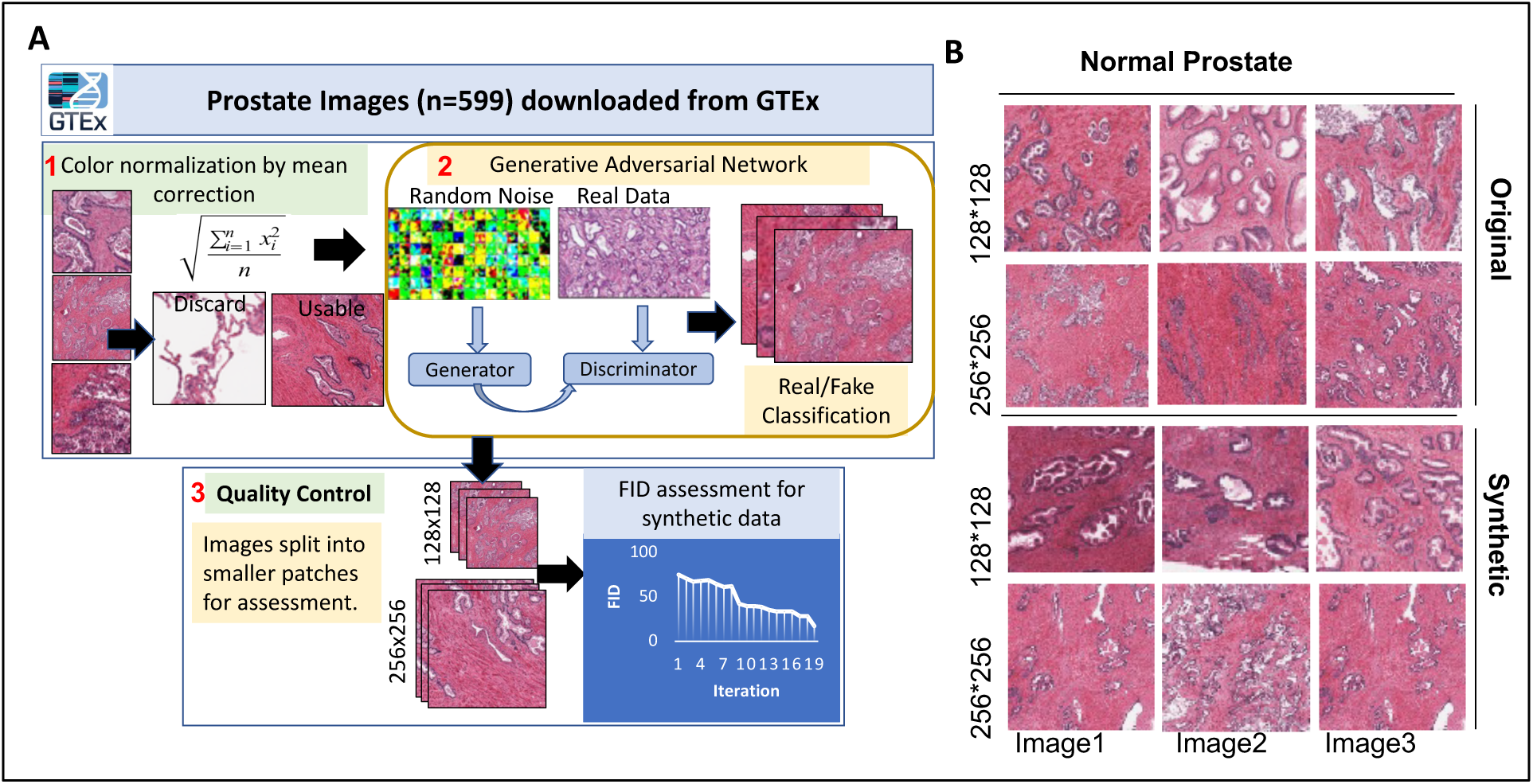
(A) Workflow for GTEX images that were used in developing the training database to be used in the GAN. Images were normalized, then fed into the GAN, and then assessed for quality. (B) Example original and synthetic histology images generated for normal prostate.

For needle biopsies, a training repository was curated, encompassing both tumor and benign sections extracted from Radbound University Medical Center and Karolinska Institute needle biopsy database. A pathologist annotated 300 needle biopsies, from which cancer-specific regions were segmented into 256×256 patches (n=1712 patches) to formulate the training repository. For benign tissues, 50 comprehensive tissue samples—including surrounding and stromal tissues were procured, adopting an identical patch size, resulting in 539 patches. Due to the inherent variances in feature dimensions between needle biopsies and RP sections, and the limited area on a needle biopsy, we investigated the effect of patch size with respect to the ability of CNN to accurately define the grade. For this, we took patch sizes range from 512*512, 256*256, 128*128, 96*96, 48*48, and 32*32, respectively. We observed an overall reduction in accuracy from 95.9% to 57.0%. Importantly, the most significant reduction was after a patch size of 64*64 (91% accurate) therefore, the input patch size for the needle biopsies was designated at 64×64 and subsequently upscaled to 256×256 to ensure model precision. To augment the training dataset, these patches underwent enhancement via a dcGAN, thus expanding the training repository to 2000 patches each for tumor and normal samples.

## Enhancing Model Performance

In an attempt to address one of the two outstanding questions in this study - determining the optimal number of synthetic images for training the EfficientNet classification model to achieve the highest possible accuracy while avoiding overfitting due to the similarity of the synthetic images - we optimized the output of the EfficientNet model using synthetic data generated by the dcGAN model. We then assessed the similarity indexing of these synthetically generated images to ensure the dcGAN model was not overfitted. The closer the similarity index is to 1, the better the alignment rate.

Synthetic images were generated from primary Gleason pattern in batches of 10K, 50K, and 100K. To evaluate the maximum unique synthetic images that could be used to train the model without compromising accuracy, we combined each image batch with the original images and subjected them to the convolutional neural network (CNN). The output images were then assessed for model overfitting using the Frechet Inception Distance (FID) score, a metric that calculates the distance between feature vectors for real and generated images. Our results indicated that, for the 10k batch size, accuracy in predicting primary pattern increased from 33% to 63% for GS3, 35% to 61% for GS4, and 41% to 68% for GS5. With the 50k batch size, accuracy increased from 33% to 70% for GS3, 35% to 65% for GS4, and 41% to 71% for GS5. For the 100k batch size, accuracy increased from 33% to 65% for GS3, 35% to 62% for GS4, and 41% to 67% for GS5. Based on these findings, we deduced that the sample complexity in the images peaked at approximately 50K and plateaued thereafter. Consequently, we incorporated a maximum of 50K synthetic images into the model.

To further mitigate the risk of overfitting the model due to minimal variation within the synthetically generated images of the same batch, we implemented a cross-validation approach to assess the inherent variability. For this purpose, all image tiles, both real and synthetic, were randomly arranged and the dataset was divided into ten subgroups. These subgroups were expected to define the training and testing sets in a randomized manner, allowing us to evaluate mean square error, misclassification error rate, and confidence intervals. Our findings, as detailed in Table 1 (left), indicate that the similarity index values ranged from 0.8 to 1.0, while the accuracy remained consistent for GS3 (0.68-0.71), GS4 (0.63-0.70), and GS5 (0.66-0.73). These results suggest that the model did not generate any significant outlier images that could confound the accuracy of the EfficientNet CNN, thus providing us with confidence in the quality and variability of the generated synthetic images.

**Table 1:** Similarity index values returned from the 10 fold cross validation run for RP images (left) and needle biopsy images (right).

| Group | Similarity Index | Group | Similarity Index |
| --- | --- | --- | --- |
| 1 | 0.8571 | 1 | 0.9281 |
| 2 | 0.9665 | 2 | 0.9663 |
| 3 | 0.9589 | 3 | 0.9219 |
| 4 | 0.8115 | 4 | 0.9354 |
| 5 | 0.9656 | 5 | 0.9451 |
| 6 | 0.9118 | 6 | 0.9846 |
| 7 | 0.9850 | 7 | 0.9372 |
| 8 | 0.8788 | 8 | 0.9558 |
| 9 | 0.9986 | 9 | 0.9992 |
| 10 | 0.8796 | 10 | 0.9891 |

Similar to the above, to mitigate the risk of overfitting the model in case of needle biopsies, we implemented a 10 fold cross-validation approach to assess the inherent variability. For this purpose, 1000 samples from the total pool of both real and synthetic, were randomly arranged and the dataset was divided into ten subgroups. Our findings, as detailed in Table 1 (right)-indicate that the similarity index values ranged from 0.92 to 1.0, while the accuracy remained consistent for GS3 (0.67-0.71), GS4 (0.63-0.70), and GS5(0.65-0.73). These results suggest that the model did not generate any significant outlier images that could confound the accuracy of the EfficientNet CNN, thus providing us with confidence in the quality and variability of the generated synthetic needle biopsy images.

## Quality Evaluation of Synthetic Images

The second question we sought to address pertains to the extent of technical variations that may exist between the synthetically generated images and the original images. This examination is crucial because each Gleason pattern possesses specific morphological characteristics that synthetically generated images should ideally replicate. For instance, Gleason pattern 3 is characterized by well-formed glands (discrete units that can be circled individually as long as they are not fused together) of varying sizes, including branching glands. These glands may be angulated or compressed, with the key feature of retaining at least a wisp of stroma between neighboring glands. If the latter is missing, then it is rated as pattern 4. The cribriform pattern is also included under pattern 4. The cribriform glands are a confluent sheet of contiguous malignant epithelial cells with multiple glandular lumina that are easily visible at low power. Gleason Pattern 5 has 2 patterns, namely comedonecrosis and cords. In comedonecrosis, central necrosis with intraluminal necrotic cells is seen within papillary/cribriform spaces, while in singular form, cells forming cords without glandular lumens are visible.

To delineate the technical variations between the original and synthetically generated images, we categorized the original images into primary Gleason pattern 3, 4, and 5. To examine individual gland formation, we subjected the images to a principal component analysis (PCA) to investigate the color distribution among GS3, GS4, and GS5. The PCA enabled us to identify features and cellular morphology inherent to each of the Gleason pattern, as described previously. Each image was represented as a large multi-layered 3D matrix, with the x and y coordinates in the matrix indicating the location on the image and the z coordinate defined by the color intensity. Three distinct numerical values were considered for color intensity, one each for red, green, and blue intensity, representing a single point or pixel on the image (Figure 3 illustrating color distribution). For Gleason pattern 3, the morphological appearance of the gland is well-formed and predominantly circular in shape. The matrices in these patterns are represented by a white color on the image and 0’s at each level of the matrices. The whitespace is encircled by darker color staining, where high-intensity values are present for the red and blue layers. Defining Gleason pattern 4 entails a change in shape that can be visually discerned by a pathologist as the well-formed structure becomes fused or shows a cribriform architecture. In the numerical matrix, the whitespace (or 0) values in the array were represented in different locations. Consequently, when examining a general PCA between Gleason pattern 3 and 4, the differences in whitespace and surrounding color wsd appear in distinct locations within the matrix and represented in the PCA as two unique groups of images (Figure 3B, C).

**Figure 3.**
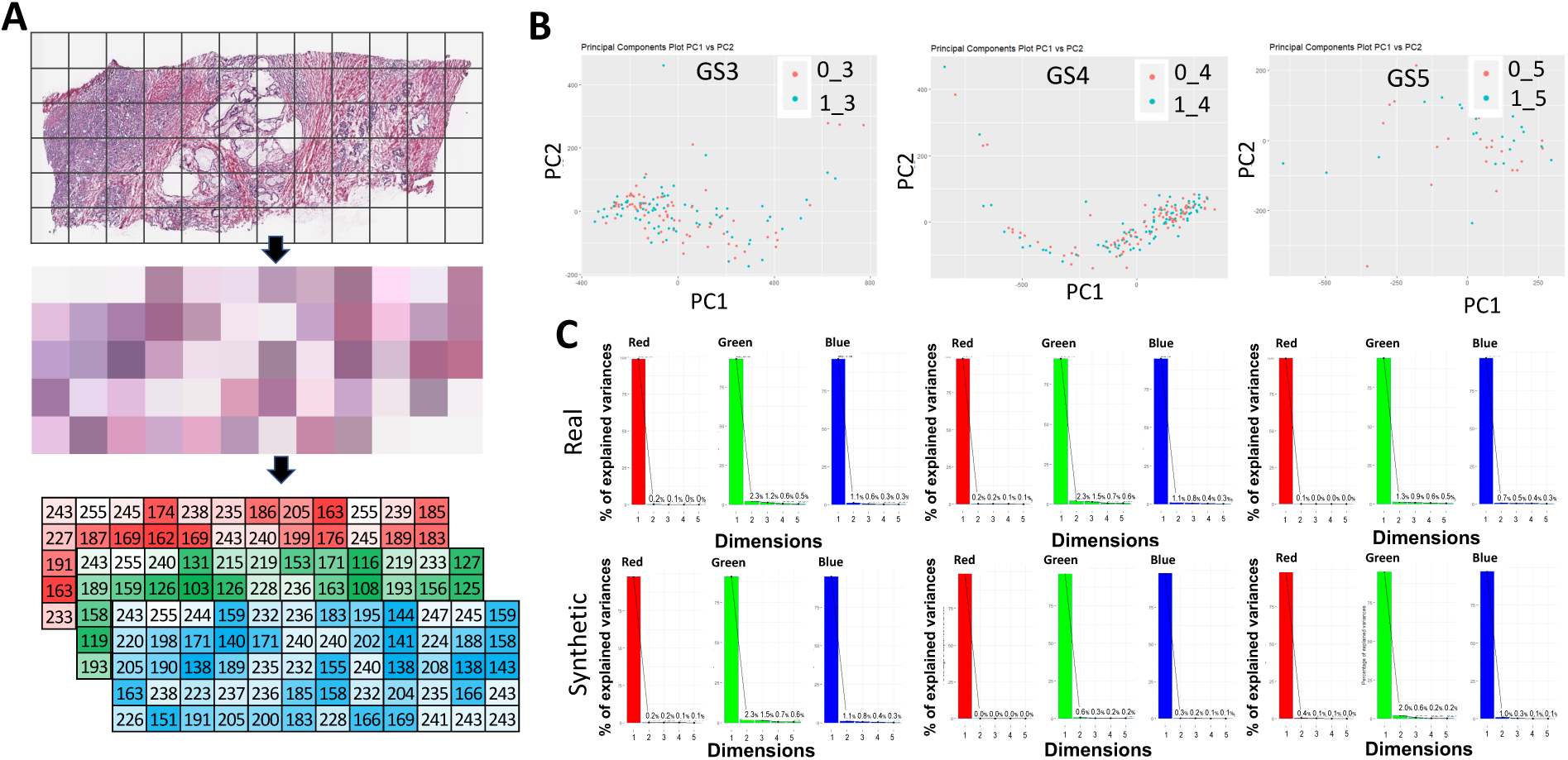
(A) Workflow how the image is chopped and then divided by RGB color matrices. (B) PCA analysis according to color gradients. Prefix of “0_” represent real images, and “1_” represent synthetic images while the suffix represents Gleason pattern. (C) Color gradient variance figures for each Gleason pattern. This represents the highest contribution of color variance to each image. *GS=Gleason pattern.

These principal considerations allowed us to employ PCA as a method for understanding the technical variations that may exist between different primary Gleason pattern in both synthetically generated and original images. In our continued investigation of technical variations, we sought to identify technical staining artifacts that could confound color variance. To achieve this, single image PCA color analysis revealed 98.8% of the variance in red, 96.1% of the variance in blue, and only 92% of the variance in green, which collectively represents all primary Gleason pattern in the original images. The least amount of variance was observed in the red and blue colors, which are the dominant colors in the tissue images. The greatest amount of variance was found in the green color, which is the least dominant color in the tissue images. The low variance results for red and blue suggest the absence of hidden confounding staining artifacts in both synthetic and original images. To determine if the level of color distribution was similar between synthetic and original images, we combined the PCA results from the three colors (red, green, and blue) for both original and synthetic images. The results revealed nonsignificant (p=0.78) technical variations when comparing the color distribution between the original (n=180) and synthetic (n=187) images. The outcomes remained consistent when comparing each primary pattern as well: Gleason 3 original versus Gleason 3 synthetic (p=0.527), Gleason 4 original versus Gleason 4 synthetic (p=0.421), and Gleason 5 original versus Gleason 5 synthetic (p=0.802), respectively. Overall, our findings suggest that the technical variability was uniform within the entire image dataset and consistent across both synthetic and original images.

## Quantification of Synthetic Images

Furthermore, to characterize and contrast the geometric structures found in both synthetic and real images, we employed Spatial Heterogeneous Recurrence Quantification Analysis (SHRQA)^100–102^, a technique that can quantitatively measure the complex microstructure based on the spatial repeating patterns. As illustrated in Figure 4, the SHRQA^102^ process comprises six pivotal steps^102^. Initially, the 2D-Discrete Wavelet Transform (2D-DWT), Haar wavelet, is executed on the image to unveil underlying patterns not readily discernible in the original form. Subsequently, each image transforms an attribute vector via the Space-Filling Curve (SFC). It is worth noting that the SFC maintains most of the spatial proximity between two pixels in the image within this attribute vector. This preservation allows for exploring the image’s geometric recurrence in vector form. A distinct trajectory representing the spatial transitions emerges by projecting the attribute vector into state space, shedding light on the image’s geometric structure properties. Applying the Quadtree method, a widely used spatial segmentation method^103,104^, the state space is segmented into exclusive subregions to extract the spatial transition patterns of the attribute vector. Then, the Iterated Function System projection converts each attribute vector into a fractal plot that captures recurrence characteristics within the fractal topology. Consequently, these fractal structures are quantified to elucidate the image’s intricate geometric properties, subsequently utilized for image profiling.

**Figure 4.**
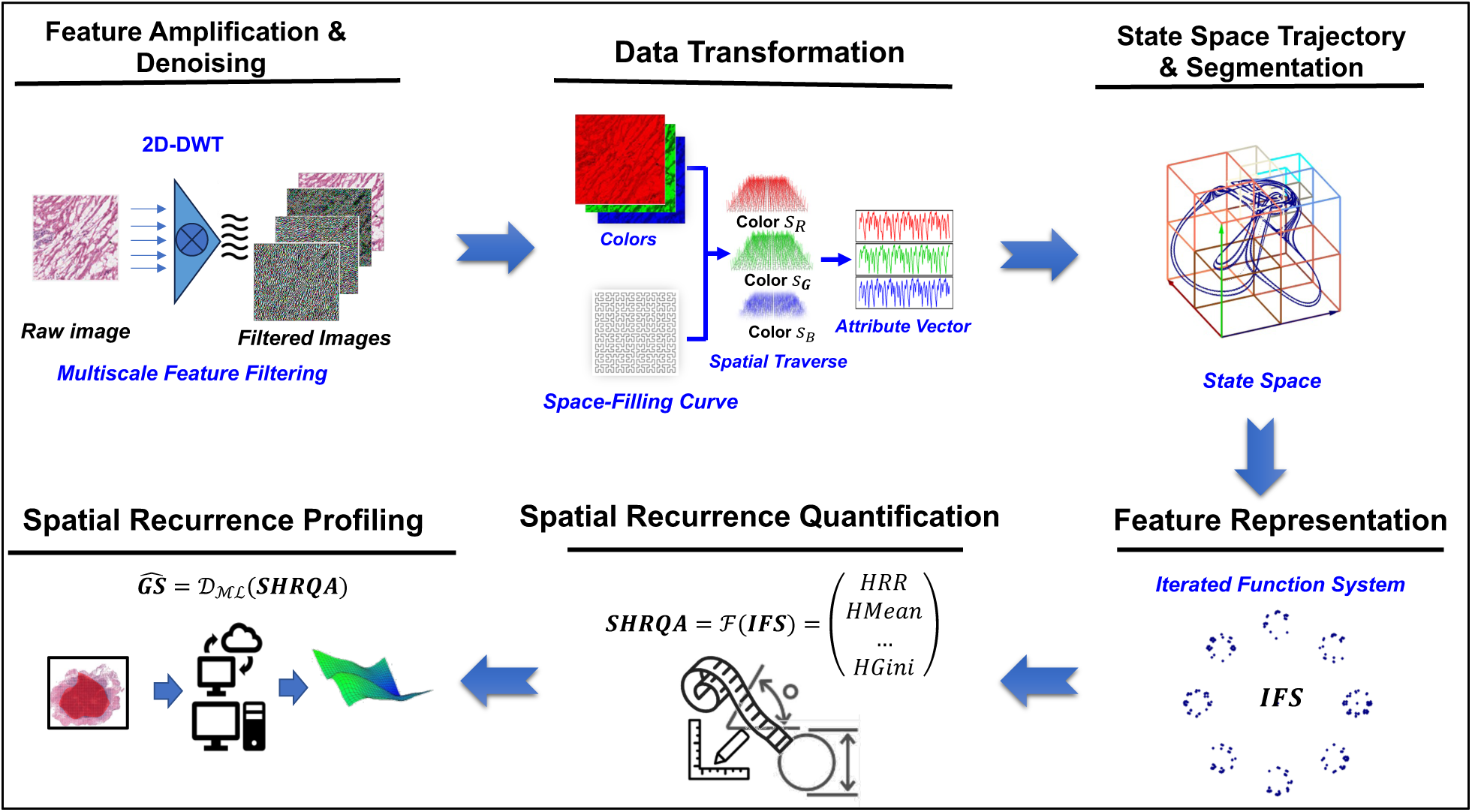
The framework of the Spatial Heterogeneous Recurrence Quantification Analysis (SHRQA). Initially, each image undergoes multiscale feature filtering using the 2D-Discrete Wavelet Transform. This amplifies intricate patterns and minimizes environmental noise. Subsequently, a space-filling curve transforms each image into an attribute vector, preserving the majority of its proximity information. Through state-space construction, pixel color/intensity transitions form a trajectory in the state space. These transitions are then projected into an Iterated Function System (IFS) to capture complex dynamic properties. The image’s nuanced geometric properties are then mathematically described using recurrence quantification analysis. Ultimately, the extracted spatial recurrence characteristics can be employed to profile images.

We employed SHRQA^102^ to analyze the spatial recurrence properties of real and synthetic image patches across three Gleason patterns (GS3/4/5) in the RP section. Half of the patches were obtained from real images, while the rest were synthetic. Notably, an even distribution of the Gleason pattern was ensured across all sample sets. We examined 3000 patches in the size 256×256 pixels, with an even split between real and synthetic patches. By applying a one-layer 2D-DWT to each image patch using Haar wavelet, we decomposed each image into four sub-images, each revealing subtle information. Subsequently, SHRQA was applied to each sub-image to quantitatively delineate each image patch’s microstructures. Initially, we extracted 1997 spatial recurrence features per patch. Following this, the Least Absolute Shrinkage and Selection Operator (LASSO) was employed to identify 1819 significant features to the Gleason pattern for further scrutiny. Subsequently, we leveraged Hotelling’s T-squared test, the multivariate counterpart of the two sample t-tests, to compare whether the spatial recurrence attributes of the real versus synthetic patches are similar. Test results indicate that the derived p-values of 0.8991 suggest no substantial disparity in spatial recurrence properties between the real and synthetic patches in the RP section. Delving into spatial recurrence properties for each Gleason pattern, the p-values of Hotelling’s T-squared tests indicate a similar conclusion that no significant difference exists between real and synthetic patches (refer to Table 2, left).

**Table 2:** Hotelling’s T-squared Two-sample Test: comparisons of spatial recurrence properties between real and synthetic images under different Gleason patterns for radical prostatectomy (RP: left) and needle biopsies (NB: right). Test results indicate that there is no significant difference between real and synthetic images in both RP and NB.

| I. Real vs. Synthetic (RP) |  |  | II. Real vs. Synthetic (NB) |  |  |
| --- | --- | --- | --- | --- | --- |
| Group | T-squared | p-value | Group | T-squared | p-value |
| All | 0.48763 | 0.8991 | All patches | 0.9634 | 0.4631 |
| GS3 | 1.0686 | 0.3837 | GS3 | 1.2786 | 0.2530 |
| GS4 | 0.51538 | 0.8802 | GS4 | 1.0291 | 0.4133 |
| GS5 | 0.51538 | 0.8802 | GS5 | 0.3984 | 0.9213 |

To illustrate the spatial recurrence properties’ similarities across different Gleason patterns in both real and synthetic patches, we employed principal component analysis (PCA) on the extracted spatial recurrence properties for RP. These results were then visualized using radar charts. Our analysis revealed that the leading ten principal components (PCs) can capture 90% of the variability in the spatial recurrence. Leveraging these PCs, we mapped the spatial property distributions of real and synthetic images across the various Gleason patterns. As depicted in Figure 5, while the distributions of spatial properties are closely aligned between real and synthetic images under the same Gleason pattern, they markedly differ when comparing different Gleason patterns.

**Figure 5.**
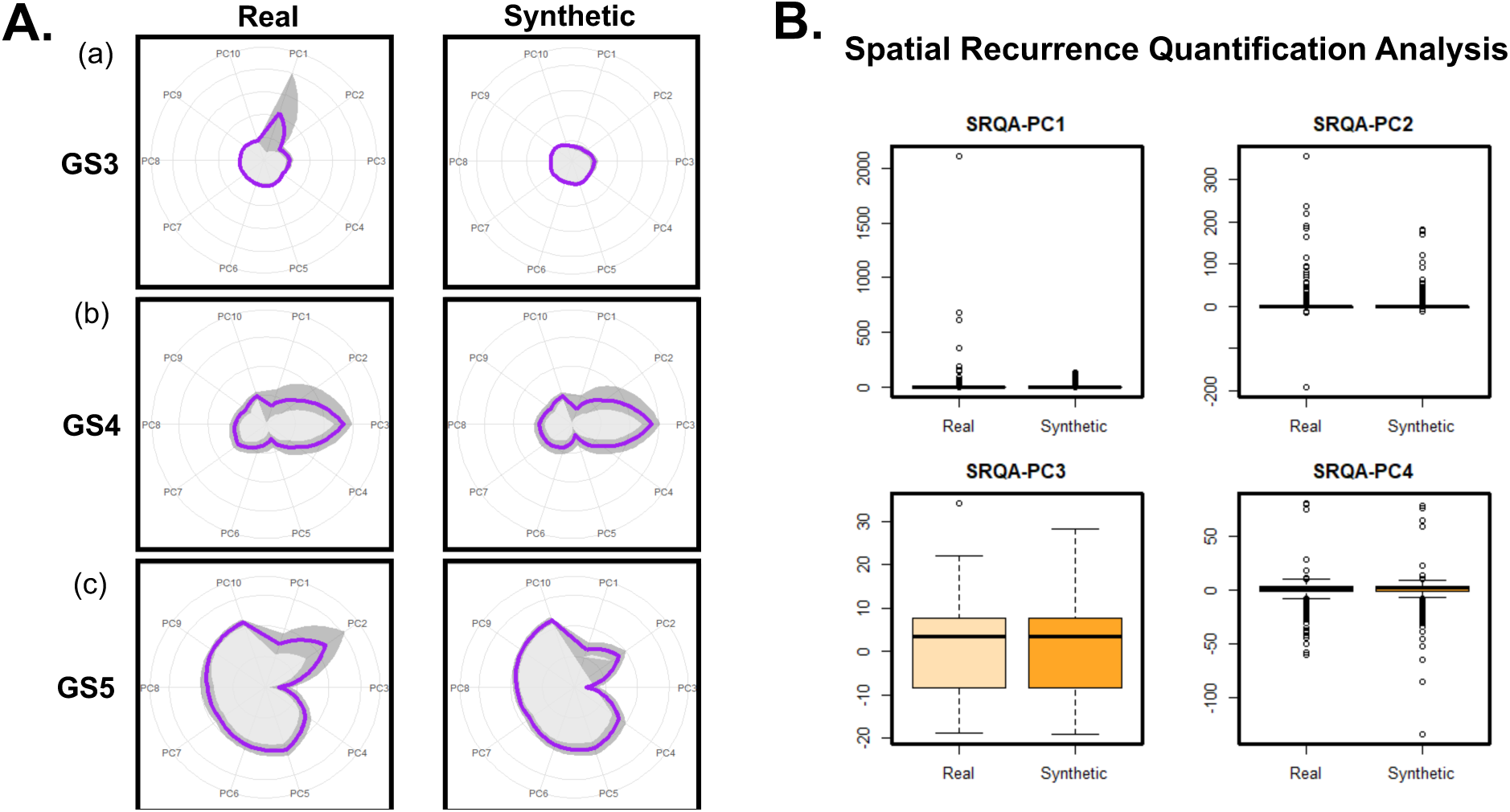
(A) The distributions of spatial recurrence properties 961 (in the first 10 Principal Components (PCs), which contain 95% of data variability) underlying different Gleason patterns for both real and synthetic patches on Radical Prostatectomy. Note that the purple lines indicate the mean values of each feature, and the gray area shows the 95% confidence interval. Our results indicate that while the distributions of spatial properties are closely aligned between real and synthetic images under the same Gleason Pattern, they markedly differ when comparing different Gleason Patterns. (B) The comparison of spatial recurrence properties between real and synthetic on the first four PCs (contain 90% of data variability). The distributions of this four PCs are similar between real and synthetic.

In addition, we analyzed the spatial recurrence properties for the Needle Biopsy (NB) section. Using the same SHRQA procedure, we examined 1200 patches of 256×256 pixels. Of these, 600 were real, and 600 were synthetic, with an even distribution across GS3, 4, and 5. A total of 2585 spatial recurrence features were extracted through SHRQA initially, and 1578 were then identified as significant features to the Gleason pattern by LASSO. The results of Hotelling’s t-squared tests (shown in Table 2, right) echo the outcomes from the RP section: synthetic images capture geometric nuances comparable to real images for any given Gleason pattern. As shown in Figure 6, for the patches in the NB section, the spatial properties’ distributions align closely between real and synthetic images with the same Gleason pattern, mirroring the findings from the RP section. Together, our results confirm that synthetic images capture geometric nuances comparable to real images for any given Gleason pattern.

**Figure 6.**
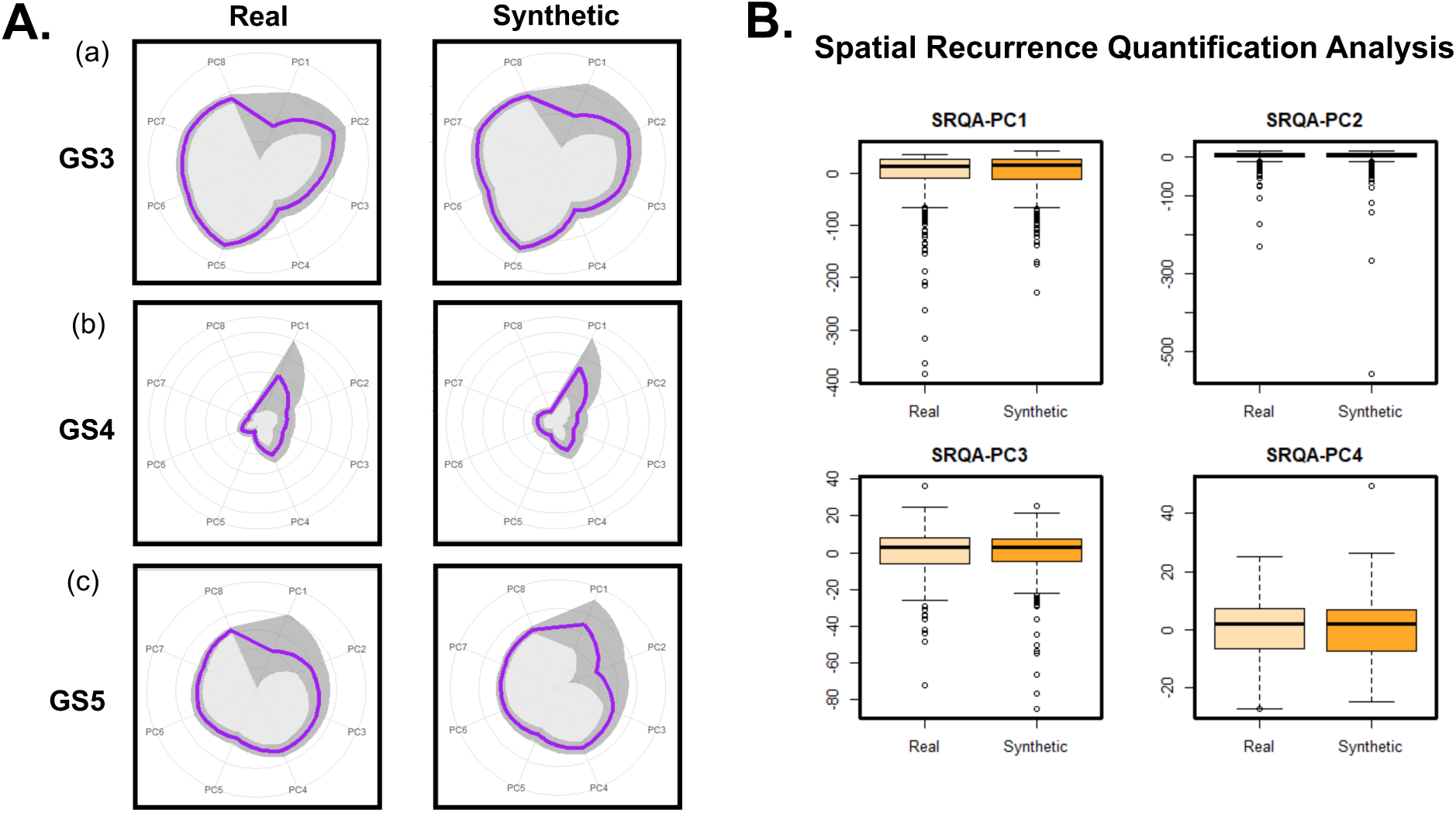
(A) The distributions of spatial recurrence properties (in the first 8 Principal Components (PCs), which contain 90% of data variability) underlying different Gleason patterns for both real and synthetic patches on Needle Biopsy. Note that the purple lines indicate the mean values of each feature, and the gray area shows the 95% confidence interval. Our results indicate that while the distributions of spatial properties are closely aligned between real and synthetic images under the same Gleason Pattern, they markedly differ when comparing different Gleason Patterns. (B) The comparison of spatial recurrence properties between real and synthetic on the first four PCs (contain 84% of data variability). The distributions of this four PCs are similar between real and synthetic.

## Validation of CNN Performance Post-Training with Enhanced Synthetic Data

Subsequently, we sought to determine whether GAN-enhanced synthetic images could substitute the RP and needle biopsy images for training a CNN model and potentially improving its grading capabilities. For this purpose, we trained the CNN model with two sources of image patches. The first source was the patches derived from original RP sections from TCGA and inhouse images, which were classified according to the Gleason pattern (normal (n=175), GS3 (n=726), GS4 (n=1029), and GS5 (n=152)). The second source of patches was derived from original digital pathology combined with the synthetically generated images, classified according to the Gleason pattern. In the second source, a total of 5000 image patches were used for each Gleason pattern. The grading capabilities of this CNN model were then compared by allowing it to assign Gleason scoring to the RP images in the TCGA (n=475). Interestingly, the results demonstrated that the CNN’s accuracy improved in GS3 from 0.53 to 0.67 (p=0.0010), in GS4 from 0.55 to 0.63 (p=0.0274), and in GS5 from 0.57 to 0.75 (p<0.0001) when trained with the combination of original and synthetic image patches compared to just being trained with original image patches. Moreover, the comparative analysis revealed a notable enhancement in accuracy and the receiver operating characteristic curve (ROC) for the combined (original and synthetic) dataset relative to the original (p= 0.0381). Additionally, the in-house RP images (n=24) were subjected to grading using CNN models described above. Results demonstrated a nonsignificant but improvement of the CNN model to accurately assign the grade from GS6 from 0.53 to 0.67 (p=0.0010), in GS7 from 0.55 to 0.63 (p=0.0274), and in GS8 from 0.57 to 0.75 (p<0.0001)

Furthermore, we extended the validation of the CNN model using needle biopsies. Similar to the RP sections, we trained the CNN model with two sources of image patches. The first source was the patches derived from original needle biopsy sections, which were classified according to the Gleason pattern (normal (n=539), GS3 (n=610), GS4 (n=890), and GS5 (n=212)). The second source of patches was derived from original digital pathology combined with the synthetically generated images, classified according to the Gleason pattern. In the second source, a total of 2000 image patches were used for each Gleason pattern. The grading capabilities of this CNN model were then compared by allowing it to assign Gleason grading to the needle biopsy images (n=3649). The comparative analysis revealed a notable enhancement in accuracy and the ROC for the amalgamated dataset relative to the original. Specifically, the enhancement was consistent across both benign and malignant samples, with an overall accuracy increase from 91% when using the original training images to an accuracy of 95% when using original and synthetic images combined (p = 0.0402). The original and synthetic combined training database yielded a sensitivity of 0.81, and a specificity of 0.92.

## DISCUSSION

In the realm of digital imagery, numerous image analysis algorithms have been developed and investigated for grading, classification, and identification of metastases across a variety of cancer types ^105–114^. Each of these models possesses unique strengths and limitations. For instance, while models such as PyTorch and TensorFlow are relatively simple to implement, they tend to have slower processing times ^115^. Another crucial factor affecting model accuracy is the architecture of the neural networks themselves. Generally, a greater number of layers in the algorithm results in higher accuracy, which also leads to slower performance ^116^. Recent studies have illustrated the potential of AI models to distinguish between non-cancerous and cancerous regions with considerable reliability ^117^. Nonetheless, most AI tools currently available face limitations in terms of their consistent application in daily cancer diagnosis and prognosis. These constraints involve the inability of existing AI tools to accurately characterize tumor heterogeneity, particularly in the context of prognosis^85^. This could be due to limitations in any one or more aspects, such as (A) Data bias: If an AI model is trained predominantly on data from one demographic patient, it may not accurately diagnose patients belonging to a different demographic background^118^. (B) Complexity of Cancer: certain cancers have a highly variable genetic makeup, making it challenging for AI to predict outcomes consistently since the AI may not have access to or be trained on such diverse genetic information^119^. (C) Technological Limitations: An AI system’s performance can degrade over time as the statistical patterns in the data it was trained on become less representative of the current clinical reality, a phenomenon known as model drift ^120^. Together, these limitations necessitate the characterization of proximal cancer areas with minimal differences in tissue architecture. In this regard, the available AI tools are substantially limited ^85^.

One of the primary reasons for these limitations is the heavy reliance of contemporary AI models on large, diverse, and unbiased clinical datasets. Such datasets are frequently scarce and challenging to access. Consequently, there is an immediate need to address these constraints to enhance the performance and applicability of AI tools in cancer diagnosis and prognosis. The findings of this study leverage machine learning and digital imagery, specifically through the use of generative adversarial network (GAN) to overcome the challenges associated with timely diagnosis and treatment. The study’s results show that an AI model trained with synthetic images competed extremely well with the model trained with original digital histology images, underscoring this novel approach’s potential. The validation of the AI model using data from the University of Miami pathology core and needle biopsy data from external sources further supports the potential generalizability and applicability of this methodology in diverse clinical settings.

Despite the promising results, there are some limitations and future directions to consider. First, further research is needed to validate the performance of the AI model in diverse patient populations, as this would provide a more comprehensive understanding of the generalizability of the findings. Second, the development of more advanced GAN models could improve the quality and diversity of synthetic images, potentially resulting in even better performance of AI models in grading prostate cancer specimens. Lastly, additional studies are needed to explore the potential of this approach in other disease contexts, including other types of cancer and non-oncological diseases.

In conclusion, this study provides strong evidence for the potential of customized GAN models in generating synthetic data that can effectively train AI models for grading prostate cancer specimens. Additionally, the use of customized GAN models for generating synthetic data could be applied to other types of cancer or diseases, potentially transforming the way AI models are trained for various medical applications. Furthermore, this methodology could be used to address the challenge of data privacy and security, as synthetic data generation can minimize the need for sharing sensitive patient information.

## CONCLUSION

This study represents the first instance of successfully generating and employing synthetically generated digital pathology data from prostate cancer to effectively train machine learning models, which subsequently demonstrated Gleason grading capabilities comparable to those trained with original patient data. While not without limitations, this research serves as an initial step toward circumventing the challenges associated with clinical data and enhancing the effectiveness of machine learning models. The models proposed in this study hold promise for improving reproducibility, reducing variability, and facilitating diagnostic and prognostic processes in prostate cancer management.

## METHODS

### Scanning and image annotation

Images are deconstructed by first taking individual areas in patchwise fashion using the software package PYHist. Briefly, the image is taken from raw .svs format and scanned for areas that are defined as tissue regions. This is done by color definition compared to whitespace background. Each predefined block is taken at a 512×512 pixel area which has been identified as the lowest amount of space that our pathologist graded any given image. These patches were then used as the training database from which other images are annotated. The image annotation for a given test image then is a whole image from which a sliding window approach is taken that moves through each of the given pixel windows and scored against the training database.

### Cohorts used

Samples were taken from TCGA. Histology images from 500 individuals were considered. 32 local samples were taken from the University of Miami pathology core. The Institutional Review Board (IRB) protocol was approved by the University of Miami Miller School of Medicine, Miami, FL, to ensure that the research adhered to ethical guidelines and principles. Furthermore, the study included 3,949 needle biopsies sourced from Radboud University Medical Center and Karolinska Institute (the PANDA challenge) ^99,121^.

### Algorithm design training and testing procedure

We selected 3 initial algorithms from which to test out the convolutional network. These were defined as the major stepwise breakthroughs in AI with the AlexNET, ResNet, and Xception models. Initially, training images were taken from a single pathologist defined areas and 10 random images were used as test images. We used granular level annotation (detail in result section) to select the model with the highest accuracy. Post model selection, the hyperparameters were turned. Here a Tree-structured Parzen Estimator was implemented to complete a sequential model optimization. In addition, tree weights were investigated by a population based training approach. From this, the highest performing hyperparameters were used along with the appropriate tree weights that were used to define the network.

### Calculation of performance

We defined accuracy in 2 different ways. First, in granular accuracy we started with the areas that were selected and annotated with the exact same Gleason pattern by all the 3 pathologists. Gold standard granular accuracy is when the convolutional network overlaps with the agreement between all pathologists. This is defined by the number of overlapping pixels in the defined area compared with the definition between the annotated areas of the pathologist pattern. Second, patient level accuracy is defined when the artificial intelligence pipeline has correctly identified a patient’s Gleason Score as defined by the TCGA pathology confirmed score.

### Development and Evaluation of a Conditional Generative Adversarial Network

A preliminary conditional Generative Adversarial Network (cGAN) was designed and implemented to assess the performance accuracy of various GAN architectures. The cGAN was developed utilizing Python 3.7.3 and the Tensorflow Keras 2.7.0 package. The generator component of the cGAN comprises three input layers and a single output layer. In parallel, the discriminator component is configured with analogous input, hidden, and output layers. The cGAN’s total parameter count was 19.2 million for each of the evaluated Gleason patterns (Supplementary Figure 1).

### Implementation and Adaptation of StyleGAN for Tissue Image Analysis

StyleGAN, a progressive generative adversarial network architecture, serves as a baseline for comparison to the cGAN, featuring a distinct generator configuration. The architecture was adopted from the original StyleGAN publication with minimal alterations to the generator and discriminator networks. A notable modification involved substituting human face images in the StyleGAN with tissue images to create a tissue image GAN. The generator’s total parameter count amounted to 28.5 million, in contrast to 26.2 million in the original StyleGAN publication and 23.1 million in a conventional generator. Particular emphasis was placed on refining the GS4 and GS5 images to ensure adequate representation of tumor heterogeneity.

### dcGAN

The dcGAN weights were initialized randomly from a normal distribution with mean = 0 and a standard deviation of 0.02. The generator neural network was constructed using 13 layers consisting of transpose function with batch normalization and ReLU functions. The walkthrough of the generator layers is as follows : ConvTranspose2d -> BatchNorm2d -> ReLU -> ConvTranspose2d -> BatchNorm2d -> ReLU -> ConvTranspose2d -> BatchNorm2d -> ReLU -> ConvTranspose2d -> BatchNorm2d -> ReLU -> ConvTrnaspose2d -> Tanh The discriminator neural network was constructed using 12 layers of Conv2d, LeakyReLU functions with batch normalization. The walkthrough of the layers is as follows : Conv2d -> LeakyReLU -> Conv2d -> BatchNorm2d -> LeakyReLU -> Con v2d -> BatchNorm2d -> LeakyReLU -> Conv2d -> BatchNorm2d -> LeakyReLU -> Conv2d -> Sigmoid Estimation of the number of parameters used for a single run ranged from 1.3T to 1.4T calculations per run to generate 1,000 synthetic images that were run on standard GPU chips (GET INFO). Time taken for a single run was dependent on the number of GPU processors available. For multi-threaded GPU the time taken ranged from 1.5 hours to 5 hours for a run while for a single GPU it ranged from 3 hours to 12 hours. Time was dependent on the level of resolution desired.

Finally, the BCE loss function was used and the Adam optimizer was implemented for both the generator and discriminator. Iteration level statistics were generated at the end of each run and saved for further analysis in matplot.

### EfficientNET

The EfficientNET baseline model was setup with the B3 function within the TensorFlow python backend. Initial testing for the B6 function was found to be too restrictive and B1 was too simple to form complex patterns so B3 was selected as our model function. To form the model based on our images, a GlobalMaxPooling2D layer was added after the initial base as well as a Dropout layer which will help avoid overfitting. The dropout rate was set to 0.2 after initial estimations were too high. The number of classes for prediction layer was set to 4 which represented the normal tissue group as compared to the primary Gleason patterns GS3, GS4, and GS5. The EfficientNET model was setup using pre-trained weights from the “imageNet” to take advantage of transfer learning to reduce analysis time.

Image augmentation was completed using Keras ImageDataGenerator function. Image rotation range was set to 45, width_shift_range and height_shift_range was set to 0.2, and the horizontal flip was set to true for flipping the image. Fill mode was defaulting to “nearest”. Validation data was not augmented, but the images were put through a rescale function to ensure that every test image was uniform before annotation.

### Statistical Calculations

FID was implemented in custom scripts developed in house. The FID model was pre-trained using Inception V3 weights for transfer learning. In house code was centered around the FID model and inserted into the dcGAN to be run during each iteration. Stats were reported at intervals of 1000 and graphed with in house python scripts.

PCA analysis was performed by first transforming the images into numerical arrays. Images were separated into normal and synthetic batches and then distributed by primary Gleason score. Intensity was calculated (using the R package imgpalr and magick) as the average of the color of the entire image while keeping the matrix framework (ie positional arguments were retained). PCA was conducted using the general prcomp function in R and plotted results were displayed in ggplot2.

### Data Sharing

De-identified participant data will be made available when all primary and secondary endpoints have been met. Any requests for trial data and supporting material (data dictionary, protocol, and statistical analysis plan) will be reviewed by the trial management group in the first instance. Only requests that have a methodologically sound proposal and whose proposed use of the data has been approved by the independent trial steering committee will be considered. Proposals should be directed to the corresponding author in the first instance; to gain access, data requestors will need to sign a data access agreement.

## Acknowledgments

Primary financial support was from Sylvester Cancer Center, the University of Miami, & Desai, and Sethi Institute of Urology. We thank all the patients whose willingness to participate made this study possible. We are grateful to the all principal investigators, sub-investigators, and local centre staff for their dedication and commitment to recruiting patients to the study. We thank members of the steering committee and data monitoring and ethics committee. We thank Dr Dipen Parekh for his valuable input from a patient perspective. The support of The Cancer Genome Atlas was essential to the successful running of the study; we thank all their staff who have contributed, past and present. We are very grateful to the laboratory team for their contribution to the study.

## Author Contributions

HA was chief investigator. DVB, and HA, designed the study and developed the protocol. OK, FI, SM, AN, VS, AN carried out the pathological analysis plan. HA and DVB coordinated the central modeling investigations. CBC developed Spatial quantification methods. CBC and DVB validated the quantifications. HA, OK and SP coordinated the data collection. DVB and HA interpreted the data. HA and DVB developed the first drafts of the manuscript. CBC, OK, SP, SM, VS, AN, FI, AB have accessed and verified all the data in the study. All authors had access to all the data reported in the study. All authors contributed to the review and amendments of the manuscript for important intellectual content and approved this final version for submission. The corresponding author had full access to all the data in the study and had final responsibility for the decision to submit for publication.

## Competiting Interests

Authors declare no compiteting interest.

## DISCLOSURES

Disclosure of Patent Information: The authors wish to inform that the technology presented in this study is part of a provisional patent application that has been filed with the United States Patent and Trademark Office (USPTO). The application has been assigned Serial No. 63/598,207 and was filed on November 13, 2023. The patent application is currently pending. Some of the authors of this paper are listed as inventors in the patent application. This patent filing may constitute a potential conflict of interest, and this statement serves to disclose this relationship in the interest of full transparency.

## CONSENT FOR PUBLICATION

All authors have provided their consent for publication.

## Notes

### Competing Interest Statement

The authors have declared no competing interest.

## REFERENCES

1 Brawley, O. W. Prostate cancer epidemiology in the United States. World J Urol 30, 195–200 (2012). 10.1007/s00345-012-0824-2

2 Badalament, R. A. & Drago, J. R. Prostate cancer. Dis Mon 37, 199–268 (1991). 10.1016/0011-5029(91)90004-u

3 Carthon, B., Sibold, H. C., Blee, S. & R, D. P. Prostate Cancer: Community Education and Disparities in Diagnosis and Treatment. Oncologist 26, 537–548 (2021). 10.1002/onco.13749

4 Cook, E. D. & Nelson, A. C. Prostate cancer screening. Curr Oncol Rep 13, 57–62 (2011). 10.1007/s11912-010-0136-x

5 Litwin, M. S. & Tan, H. J. The Diagnosis and Treatment of Prostate Cancer: A Review. Jama 317, 2532–2542 (2017). 10.1001/jama.2017.7248

6 Moon, T. D. Prostate cancer. J Am Geriatr Soc 40, 622–627 (1992). 10.1111/j.1532-5415.1992.tb02116.x

7 Schatten, H. Brief Overview of Prostate Cancer Statistics, Grading, Diagnosis and Treatment Strategies. Adv Exp Med Biol 1095, 1–14 (2018). 10.1007/978-3-319-95693-0_1

8 Wozniak-Petrofsky, J. The significance of prostatic specific antigen in men with prostate disease. An elevated PSA level may indicate prostate cancer--and it may not. Geriatr Nurs 14, 150–151 (1993). 10.1016/s0197-4572(06)80133-3

9 Aghdam, A. M. et al. MicroRNAs as Diagnostic, Prognostic, and Therapeutic Biomarkers in Prostate Cancer. Crit Rev Eukaryot Gene Expr 29, 127–139 (2019). 10.1615/CritRevEukaryotGeneExpr.2019025273

10 Albertsen, P. C. PSA testing, cancer treatment, and prostate cancer mortality reduction: What is the mechanism? Urol Oncol 41, 78–81 (2023). 10.1016/j.urolonc.2021.08.010

11 Barry, M. J. & Simmons, L. H. Prevention of Prostate Cancer Morbidity and Mortality: Primary Prevention and Early Detection. Med Clin North Am 101, 787–806 (2017). 10.1016/j.mcna.2017.03.009

12 Borley, N. & Feneley, M. R. Prostate cancer: diagnosis and staging. Asian J Androl 11, 74–80 (2009). 10.1038/aja.2008.19

13 Jalloh, M. & Cooperberg, M. R. Implementation of PSA-based active surveillance in prostate cancer. Biomark Med 8, 747–753 (2014). 10.2217/bmm.14.5

14 Merriel, S. W. D., Funston, G. & Hamilton, W. Prostate Cancer in Primary Care. Adv Ther 35, 1285–1294 (2018). 10.1007/s12325-018-0766-1

15 Nguyen-Nielsen, M. & Borre, M. Diagnostic and Therapeutic Strategies for Prostate Cancer. Semin Nucl Med 46, 484–490 (2016). 10.1053/j.semnuclmed.2016.07.002

16 Pezaro, C., Woo, H. H. & Davis, I. D. Prostate cancer: measuring PSA. Intern Med J 44, 433–440 (2014). 10.1111/imj.12407

17 Sharma, S., Zapatero-Rodríguez, J. & O’Kennedy, R. Prostate cancer diagnostics: Clinical challenges and the ongoing need for disruptive and effective diagnostic tools. Biotechnol Adv 35, 135–149 (2017). 10.1016/j.biotechadv.2016.11.009

18 Troyer, D. A., Mubiru, J., Leach, R. J. & Naylor, S. L. Promise and challenge: Markers of prostate cancer detection, diagnosis and prognosis. Dis Markers 20, 117–128 (2004). 10.1155/2004/509276

19 Venderbos, L. D. & Roobol, M. J. PSA-based prostate cancer screening: the role of active surveillance and informed and shared decision making. Asian J Androl 13, 219–224 (2011). 10.1038/aja.2010.180

20 Parekh, D. J. et al. A multi-institutional prospective trial in the USA confirms that the 4Kscore accurately identifies men with high-grade prostate cancer. Eur Urol 68, 464–470 (2015). 10.1016/j.eururo.2014.10.021

21 Borque-Fernando, Á. et al. Role of the 4Kscore test as a predictor of reclassification in prostate cancer active surveillance. Prostate Cancer Prostatic Dis 22, 84–90 (2019). 10.1038/s41391-018-0074-5

22 Konety, B. et al. The 4Kscore® Test Reduces Prostate Biopsy Rates in Community and Academic Urology Practices. Rev Urol 17, 231–240 (2015).

23 Mi, C., Bai, L., Yang, Y., Duan, J. & Gao, L. 4Kscore diagnostic value in patients with high-grade prostate cancer using cutoff values of 7.5% to 10%: A meta-analysis. Urol Oncol 39, 366.e361–366.e310 (2021). 10.1016/j.urolonc.2020.11.001

24 Punnen, S. et al. The 4Kscore Predicts the Grade and Stage of Prostate Cancer in the Radical Prostatectomy Specimen: Results from a Multi-institutional Prospective Trial. Eur Urol Focus 3, 94–99 (2017). 10.1016/j.euf.2015.12.005

25 Scuderi, S. et al. Implementation of 4Kscore as a Secondary Test Before Prostate Biopsy: Impact on US Population Trends for Prostate Cancer. Eur Urol Open Sci 52, 1–3 (2023). 10.1016/j.euros.2023.03.011

26 Zappala, S. M. et al. The 4Kscore blood test accurately identifies men with aggressive prostate cancer prior to prostate biopsy with or without DRE information. Int J Clin Pract 71 (2017). 10.1111/ijcp.12943

27 Hessels, D. et al. DD3(PCA3)-based molecular urine analysis for the diagnosis of prostate cancer. Eur Urol 44, 8–15; discussion 15-16 (2003). 10.1016/s0302-2838(03)00201-x

28 Day, J. R., Jost, M., Reynolds, M. A., Groskopf, J. & Rittenhouse, H. PCA3: from basic molecular science to the clinical lab. Cancer Lett 301, 1–6 (2011). 10.1016/j.canlet.2010.10.019

29 de la Taille, A. Progensa PCA3 test for prostate cancer detection. Expert Rev Mol Diagn 7, 491–497 (2007). 10.1586/14737159.7.5.491

30 Filella, X. et al. PCA3 in the detection and management of early prostate cancer. Tumour Biol 34, 1337–1347 (2013). 10.1007/s13277-013-0739-6

31 Gunelli, R., Fragalà, E. & Fiori, M. PCA3 in Prostate Cancer. Methods Mol Biol 2292, 105–113 (2021). 10.1007/978-1-0716-1354-2_9

32 Hessels, D. & Schalken, J. A. The use of PCA3 in the diagnosis of prostate cancer. Nat Rev Urol 6, 255–261 (2009). 10.1038/nrurol.2009.40

33 Yang, Z., Yu, L. & Wang, Z. PCA3 and TMPRSS2-ERG gene fusions as diagnostic biomarkers for prostate cancer. Chin J Cancer Res 28, 65–71 (2016). 10.3978/j.issn.1000-9604.2016.01.05

34 Kim, L. et al. Clinical utility and cost modelling of the phi test to triage referrals into image-based diagnostic services for suspected prostate cancer: the PRIM (Phi to RefIne Mri) study. BMC Med 18, 95 (2020). 10.1186/s12916-020-01548-3

35 Barisiene, M. et al. Prostate Health Index and Prostate Health Index Density as Diagnostic Tools for Improved Prostate Cancer Detection. Biomed Res Int 2020, 9872146 (2020). 10.1155/2020/9872146

36 Tosoian, J. J. et al. Use of the Prostate Health Index for detection of prostate cancer: results from a large academic practice. Prostate Cancer Prostatic Dis 20, 228–233 (2017). 10.1038/pcan.2016.72

37 Yan, J. Q. et al. Prostate Health Index (phi) and its derivatives predict Gleason score upgrading after radical prostatectomy among patients with low-risk prostate cancer. Asian J Androl 24, 406–410 (2022). 10.4103/aja202174

38 Ye, C. et al. The Prostate Health Index and multi-parametric MRI improve diagnostic accuracy of detecting prostate cancer in Asian populations. Investig Clin Urol 63, 631–638 (2022). 10.4111/icu.20220056

39 Zhang, G. et al. Assessment on clinical value of prostate health index in the diagnosis of prostate cancer. Cancer Med 8, 5089–5096 (2019). 10.1002/cam4.2376

40 Kaplan, I. et al. Real time MRI-ultrasound image guided stereotactic prostate biopsy. Magn Reson Imaging 20, 295–299 (2002). 10.1016/s0730-725x(02)00490-3

41 Fernandes, M. C., Yildirim, O., Woo, S., Vargas, H. A. & Hricak, H. The role of MRI in prostate cancer: current and future directions. Magma 35, 503–521 (2022). 10.1007/s10334-022-01006-6

42 Mendhiratta, N., Taneja, S. S. & Rosenkrantz, A. B. The role of MRI in prostate cancer diagnosis and management. Future Oncol 12, 2431–2443 (2016). 10.2217/fon-2016-0169

43 Stabile, A. et al. Multiparametric MRI for prostate cancer diagnosis: current status and future directions. Nat Rev Urol 17, 41–61 (2020). 10.1038/s41585-019-0212-4

44 Stempel, C. V., Dickinson, L. & Pendsé, D. MRI in the Management of Prostate Cancer. Semin Ultrasound CT MR 41, 366–372 (2020). 10.1053/j.sult.2020.04.003

45 Wibmer, A. G., Vargas, H. A. & Hricak, H. Role of MRI in the diagnosis and management of prostate cancer. Future Oncol 11, 2757–2766 (2015). 10.2217/fon.15.206

46 Ikeda, S., Elkin, S. K., Tomson, B. N., Carter, J. L. & Kurzrock, R. Next-generation sequencing of prostate cancer: genomic and pathway alterations, potential actionability patterns, and relative rate of use of clinical-grade testing. Cancer Biol Ther 20, 219–226 (2019). 10.1080/15384047.2018.1523849

47 Abdulmajed, M. I., Hughes, D. & Shergill, I. S. The role of transperineal template biopsies of the prostate in the diagnosis of prostate cancer: a review. Expert Rev Med Devices 12, 175–182 (2015). 10.1586/17434440.2015.990376

48 Ahdoot, M. et al. MRI-Targeted, Systematic, and Combined Biopsy for Prostate Cancer Diagnosis. N Engl J Med 382, 917–928 (2020). 10.1056/NEJMoa1910038

49 He, Y., Shen, Q., Fu, W., Wang, H. & Song, G. Optimized grade group for reporting prostate cancer grade in systematic and MRI-targeted biopsies. Prostate 82, 1125–1132 (2022). 10.1002/pros.24365

50 Pinto, F. et al. Imaging in prostate cancer diagnosis: present role and future perspectives. Urol Int 86, 373–382 (2011). 10.1159/000324515

51 Bangma, C. H., Roemeling, S. & Schroder, F. H. Overdiagnosis and overtreatment of early detected prostate cancer. World J Urol 25, 3–9 (2007). 10.1007/s00345-007-0145-z

52 Loeb, S. et al. Overdiagnosis and overtreatment of prostate cancer. Eur Urol 65, 1046–1055 (2014). 10.1016/j.eururo.2013.12.062

53 Thompson, I. M. Overdiagnosis and overtreatment of prostate cancer. Am Soc Clin Oncol Educ Book, e35–39 (2012). 10.14694/EdBook_AM.2012.32.98

54 Resnick, M. J. et al. Repeat prostate biopsy and the incremental risk of clinically insignificant prostate cancer. Urology 77, 548–552 (2011). 10.1016/j.urology.2010.08.063

55 Tataru, O. S. et al. Artificial Intelligence and Machine Learning in Prostate Cancer Patient Management-Current Trends and Future Perspectives. Diagnostics (Basel) 11 (2021). 10.3390/diagnostics11020354

56 Choi, R. Y., Coyner, A. S., Kalpathy-Cramer, J., Chiang, M. F. & Campbell, J. P. Introduction to Machine Learning, Neural Networks, and Deep Learning. Transl Vis Sci Technol 9, 14 (2020). 10.1167/tvst.9.2.14

57 Deo, R. C. Machine Learning in Medicine. Circulation 132, 1920–1930 (2015). 10.1161/circulationaha.115.001593

58 Lo Vercio, L., et al. Supervised machine learning tools: a tutorial for clinicians. J Neural Eng 17 (2020). 10.1088/1741-2552/abbff2

59 Villoutreix, P. What machine learning can do for developmental biology. Development 148 (2021). 10.1242/dev.188474

60 Colling, R. et al. Artificial intelligence in digital pathology: a roadmap to routine use in clinical practice. J Pathol 249, 143–150 (2019). 10.1002/path.5310

61 Yousif, M. et al. Artificial intelligence applied to breast pathology. Virchows Arch 480, 191–209 (2022). 10.1007/s00428-021-03213-3

62 Jovic, S., Miljkovic, M., Ivanovic, M., Saranovic, M. & Arsic, M. Prostate Cancer Probability Prediction By Machine Learning Technique. Cancer Invest 35, 647–651 (2017). 10.1080/07357907.2017.1406496

63 Arvaniti, E. et al. Automated Gleason grading of prostate cancer tissue microarrays via deep learning. Sci Rep 8, 12054 (2018). 10.1038/s41598-018-30535-1

64 Bhattacharya, I. et al. Bridging the gap between prostate radiology and pathology through machine learning. Med Phys 49, 5160–5181 (2022). 10.1002/mp.15777

65 Bulten, W. et al. Automated deep-learning system for Gleason grading of prostate cancer using biopsies: a diagnostic study. Lancet Oncol 21, 233–241 (2020). 10.1016/s1470-2045(19)30739-9

66 Duenweg, S. R. et al. Comparison of a machine and deep learning model for automated tumor annotation on digitized whole slide prostate cancer histology. PLoS One 18, e0278084 (2023). 10.1371/journal.pone.0278084

67 Le, M. H. et al. Automated diagnosis of prostate cancer in multi-parametric MRI based on multimodal convolutional neural networks. Phys Med Biol 62, 6497–6514 (2017). 10.1088/1361-6560/aa7731

68 Li, H. et al. Machine Learning in Prostate MRI for Prostate Cancer: Current Status and Future Opportunities. Diagnostics (Basel) 12 (2022). 10.3390/diagnostics12020289

69 Mehralivand, S. et al. Deep learning-based artificial intelligence for prostate cancer detection at biparametric MRI. Abdom Radiol (NY) 47, 1425–1434 (2022). 10.1007/s00261-022-03419-2

70 Xiang, J. et al. Automatic diagnosis and grading of Prostate Cancer with weakly supervised learning on whole slide images. Comput Biol Med 152, 106340 (2023). 10.1016/j.compbiomed.2022.106340

71 Bhargava, H. K. et al. Computationally Derived Image Signature of Stromal Morphology Is Prognostic of Prostate Cancer Recurrence Following Prostatectomy in African American Patients. Clin Cancer Res 26, 1915–1923 (2020). 10.1158/1078-0432.Ccr-19-2659

72 Safarpoor, A., Kalra, S. & Tizhoosh, H. R. Generative models in pathology: synthesis of diagnostic quality pathology images(†). J Pathol 253, 131–132 (2021). 10.1002/path.5577

73 Xu, I. R. L. et al. Generative Adversarial Networks Can Create High Quality Artificial Prostate Cancer Magnetic Resonance Images. J Pers Med 13 (2023). 10.3390/jpm13030547

74 Alfano, R. et al. Prostate cancer classification using radiomics and machine learning on mp-MRI validated using co-registered histology. Eur J Radiol 156, 110494 (2022). 10.1016/j.ejrad.2022.110494

75 Bertelli, E. et al. Machine and Deep Learning Prediction Of Prostate Cancer Aggressiveness Using Multiparametric MRI. Front Oncol 11, 802964 (2021). 10.3389/fonc.2021.802964

76 Bonekamp, D. & Schlemmer, H. P. [Machine learning and multiparametric MRI for early diagnosis of prostate cancer]. Urologe A 60, 576–591 (2021). 10.1007/s00120-021-01492-x

77 Cuocolo, R. et al. Machine learning for the identification of clinically significant prostate cancer on MRI: a meta-analysis. Eur Radiol 30, 6877–6887 (2020). 10.1007/s00330-020-07027-w

78 Cuocolo, R. et al. Machine learning applications in prostate cancer magnetic resonance imaging. Eur Radiol Exp 3, 35 (2019). 10.1186/s41747-019-0109-2

79 Hosseinzadeh, M. et al. Deep learning-assisted prostate cancer detection on bi-parametric MRI: minimum training data size requirements and effect of prior knowledge. Eur Radiol 32, 2224–2234 (2022). 10.1007/s00330-021-08320-y

80 Michaely, H. J., Aringhieri, G., Cioni, D. & Neri, E. Current Value of Biparametric Prostate MRI with Machine-Learning or Deep-Learning in the Detection, Grading, and Characterization of Prostate Cancer: A Systematic Review. Diagnostics (Basel) 12 (2022). 10.3390/diagnostics12040799

81 Vente, C., Vos, P., Hosseinzadeh, M., Pluim, J. & Veta, M. Deep Learning Regression for Prostate Cancer Detection and Grading in Bi-Parametric MRI. IEEE Trans Biomed Eng 68, 374–383 (2021). 10.1109/tbme.2020.2993528

82 Madabhushi, A. & Lee, G. Image analysis and machine learning in digital pathology: Challenges and opportunities. Med Image Anal 33, 170–175 (2016). 10.1016/j.media.2016.06.037

83 Freeman, K. et al. Use of artificial intelligence for image analysis in breast cancer screening programmes: systematic review of test accuracy. BMJ 374, n1872 (2021). 10.1136/bmj.n1872

84 Twilt, J. J., van Leeuwen, K. G., Huisman, H. J., Futterer, J. J. & de Rooij, M. Artificial Intelligence Based Algorithms for Prostate Cancer Classification and Detection on Magnetic Resonance Imaging: A Narrative Review. Diagnostics (Basel) 11 (2021). 10.3390/diagnostics11060959

85 Van Booven, D. J. et al. A Systematic Review of Artificial Intelligence in Prostate Cancer. Res Rep Urol 13, 31–39 (2021). 10.2147/RRU.S268596

86 Bulten, W. et al. Artificial intelligence assistance significantly improves Gleason grading of prostate biopsies by pathologists. Mod Pathol 34, 660–671 (2021). 10.1038/s41379-020-0640-y

87 Bulten, W. et al. Artificial intelligence for diagnosis and Gleason grading of prostate cancer: the PANDA challenge. Nat Med 28, 154–163 (2022). 10.1038/s41591-021-01620-2

88 Shah, P. et al. Artificial intelligence and machine learning in clinical development: a translational perspective. NPJ Digit Med 2, 69 (2019). 10.1038/s41746-019-0148-3

89 Weissler, E. H. et al. The role of machine learning in clinical research: transforming the future of evidence generation. Trials 22, 537 (2021). 10.1186/s13063-021-05489-x

90 Banerjee, I. et al. Weakly supervised natural language processing for assessing patient-centered outcome following prostate cancer treatment. JAMIA Open 2, 150–159 (2019). 10.1093/jamiaopen/ooy057

91 Yagi, Y. et al. Development of a database system and image viewer to assist in the correlation of histopathologic features and digital image analysis with clinical and molecular genetic information. Pathol Int 66, 63–74 (2016). 10.1111/pin.12382

92 Collet, J. P. [Limitations of clinical trials]. Rev Prat 50, 833–837 (2000).

93 Cheng, J. Y., Abel, J. T., Balis, U. G. J., McClintock, D. S. & Pantanowitz, L. Challenges in the Development, Deployment, and Regulation of Artificial Intelligence in Anatomic Pathology. Am J Pathol 191, 1684–1692 (2021). 10.1016/j.ajpath.2020.10.018

94 van der Laak, J., Litjens, G. & Ciompi, F. Deep learning in histopathology: the path to the clinic. Nat Med 27, 775–784 (2021). 10.1038/s41591-021-01343-4

95 Krzyszczyk, P. et al. The growing role of precision and personalized medicine for cancer treatment. TECHNOLOGY 06, 79–100 (2018). 10.1142/s2339547818300020

96 Zerbe, N., Hufnagl, P. & Schlüns, K. Distributed computing in image analysis using open source frameworks and application to image sharpness assessment of histological whole slide images. Diagn Pathol 6 **Suppl 1**, S16 (2011). 10.1186/1746-1596-6-s1-s16

97 Aeffner, F. et al. Digital Microscopy, Image Analysis, and Virtual Slide Repository. ILAR Journal 59, 66–79 (2018). 10.1093/ilar/ily007

98 McAlpine, E. D., Pantanowitz, L. & Michelow, P. M. Challenges Developing Deep Learning Algorithms in Cytology. Acta Cytol 65, 301–309 (2021). 10.1159/000510991

99 Bulten, W. et al. Artificial intelligence for diagnosis and Gleason grading of prostate cancer: the PANDA challenge. Nat Med 28, 154–163 (2022). 10.1038/s41591-021-01620-2

100 Chen, C. B., Wang, Y., Fu, X. & Yang, H. Recurrence Network Analysis of Histopathological Images for the Detection of Invasive Ductal Carcinoma in Breast Cancer. IEEE/ACM Trans Comput Biol Bioinform 20, 3234–3244 (2023). 10.1109/TCBB.2023.3282798

101 Chen, C. B., Yang, H. & Kumara, S. Recurrence network modeling and analysis of spatial data. Chaos 28, 085714 (2018). 10.1063/1.5024917

102 Yang, H., Chen, C. B. & Kumara, S. Heterogeneous recurrence analysis of spatial data. Chaos 30, 013119 (2020). 10.1063/1.5129959

103 Shukla, P., Verma, A., Abhishek, Verma, S. & Kumar, M. Interpreting SVM for medical images using Quadtree. Multimed Tools Appl 79, 29353–29373 (2020). 10.1007/s11042-020-09431-2

104 Baia, J., Zhao, X. & Chenb, J. Z.

105 Niazi, M. K. K. et al. Visually Meaningful Histopathological Features for Automatic Grading of Prostate Cancer. IEEE J Biomed Health Inform 21, 1027–1038 (2017). 10.1109/JBHI.2016.2565515

106 Erratum to “Visually Meaningful Histopathological Features for Automatic Grading of Prostate Cancer”. IEEE J Biomed Health Inform 21, 1473–1474 (2017). 10.1109/JBHI.2017.2733238

107 Hou, L. et al. Patch-based Convolutional Neural Network for Whole Slide Tissue Image Classification. Proc IEEE Comput Soc Conf Comput Vis Pattern Recognit 2016, 2424–2433 (2016). 10.1109/CVPR.2016.266

108 Ren, J. et al. Computer aided analysis of prostate histopathology images Gleason grading especially for Gleason score 7. Annu Int Conf IEEE Eng Med Biol Soc 2015, 3013–3016 (2015). 10.1109/EMBC.2015.7319026

109 Fauzi, M. F. et al. Classification of follicular lymphoma: the effect of computer aid on pathologists grading. BMC Med Inform Decis Mak 15, 115 (2015). 10.1186/s12911-015-0235-6

110 Kothari, S., Phan, J. H., Young, A. N. & Wang, M. D. Histological image classification using biologically interpretable shape-based features. BMC Med Imaging 13, 9 (2013). 10.1186/1471-2342-13-9

111 Kong, J. et al. Machine-based morphologic analysis of glioblastoma using whole-slide pathology images uncovers clinically relevant molecular correlates. PLoS One 8, e81049 (2013). 10.1371/journal.pone.0081049

112 Dundar, M. M. et al. Computerized classification of intraductal breast lesions using histopathological images. IEEE Trans Biomed Eng 58, 1977–1984 (2011). 10.1109/TBME.2011.2110648

113 Sertel, O. et al. Computer-aided Prognosis of Neuroblastoma on Whole-slide Images: Classification of Stromal Development. Pattern Recognit 42, 1093–1103 (2009). 10.1016/j.patcog.2008.08.027

114 Kong, J. et al. Computer-assisted grading of neuroblastic differentiation. Arch Pathol Lab Med 132, 903–904; author reply 904 (2008). 10.1043/1543-2165(2008)132[903:CGOND]2.0.CO;210.5858/2008-132-903-CGOND

115 Munjal, R., Arif, S., Wendler, F. & Kanoun, O. Comparative Study of Machine-Learning Frameworks for the Elaboration of Feed-Forward Neural Networks by Varying the Complexity of Impedimetric Datasets Synthesized Using Eddy Current Sensors for the Characterization of Bi-Metallic Coins. Sensors (Basel) 22 (2022). 10.3390/s22041312

116 Hodas, N. O. & Stinis, P. Doing the Impossible: Why Neural Networks Can Be Trained at All. Front Psychol 9, 1185 (2018). 10.3389/fpsyg.2018.01185

117 Koh, D. M. et al. Artificial intelligence and machine learning in cancer imaging. Commun Med (Lond) 2, 133 (2022). 10.1038/s43856-022-00199-0

118 Celi, L. A. et al. Sources of bias in artificial intelligence that perpetuate healthcare disparities-A global review. PLOS Digit Health 1, e0000022 (2022). 10.1371/journal.pdig.0000022

119 Shreve, J. T., Khanani, S. A. & Haddad, T. C. Artificial Intelligence in Oncology: Current Capabilities, Future Opportunities, and Ethical Considerations. Am Soc Clin Oncol Educ Book 42, 1–10 (2022). 10.1200/EDBK_350652

120 Khan, B. et al. Drawbacks of Artificial Intelligence and Their Potential Solutions in the Healthcare Sector. Biomed Mater Devices, 1–8 (2023). 10.1007/s44174-023-00063-2

121 Singhal, N. et al. A deep learning system for prostate cancer diagnosis and grading in whole slide images of core needle biopsies. Sci Rep 12, 3383 (2022). 10.1038/s41598-022-07217-0

