## Supplementary data for "Synthetic Histology Images for Training AI Models: A Novel Approach to Improve Prostate Cancer Diagnosis"

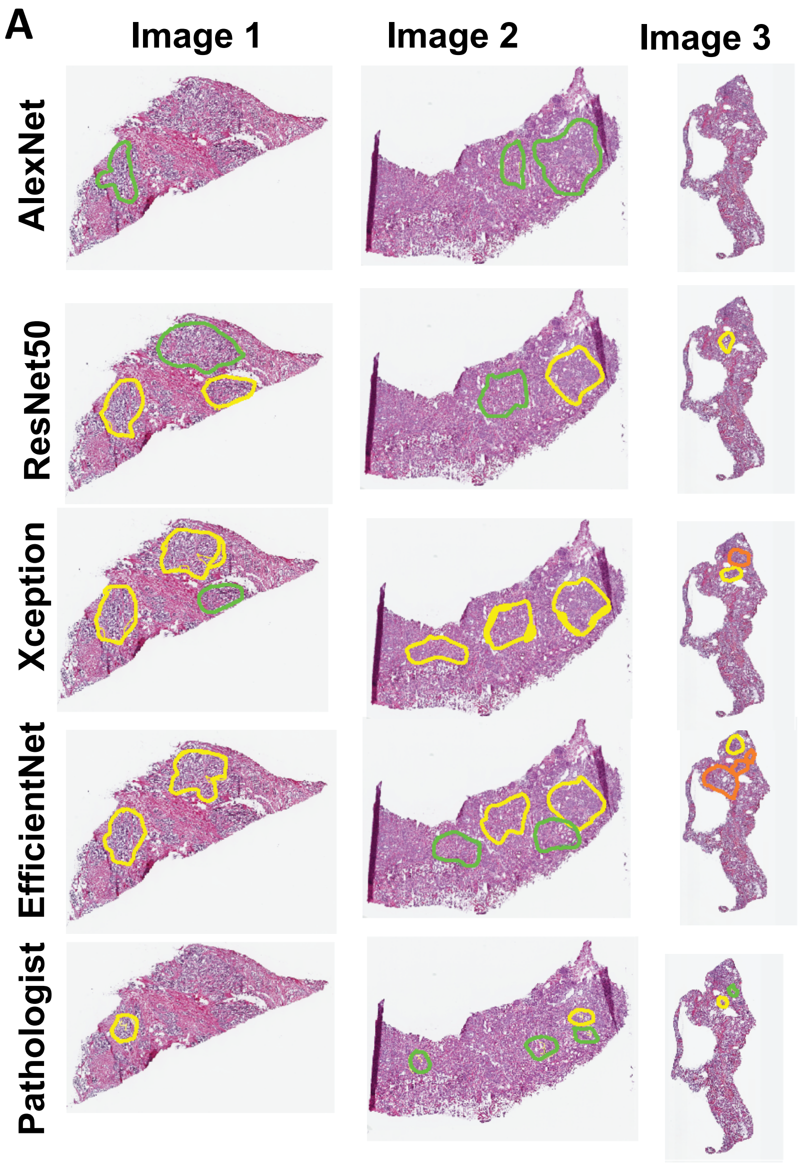


**Supplementary Figure 1.** (A) Annotation by the output from the AlexNet, ResNet50, and EfficientNet models for a given tissue image. Regions in an individual image with different Gleason patterns are distinguished using different colors (Colors from green to yellow to red for Gleason patterns 3,4,5, respectively). (B) The AI models assigned Gleason patterns.

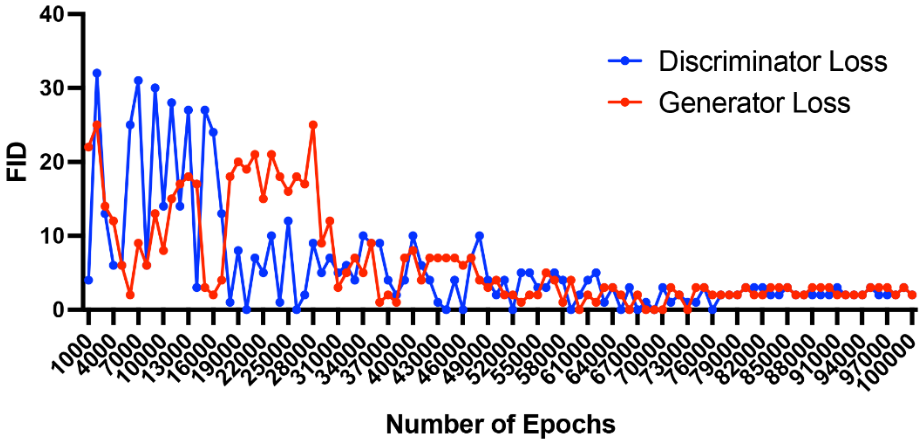


**Supplemental Figure 2**. Loss function graphs show that graphs stabilize over time and optimal epoch cutoff.

| | AlexNet | ResNet | Xception | EfficientNet |
|---|---|---|---|---|
| *Normal* | 3 | 3 | 4 | 4 |
| *3+3* | 2 | 2 | 2 | 2 |
| *3+4* | 1 | 1 | 2 | 1 |
| *4+3* | 2 | 1 | 1 | 1 |
| *4+4* | 1 | 1 | 1 | 2 |
| *4+5* | 2 | 3 | 2 | 3 |
| *# correct* | 11 | 11 | 12 | 13 |

**Supplemental Table 1**. Outcomes of image analysis by CNN models such as AlexNet, ResNet, Xception, and EfficientNet evaluation.
